# Integrated single-cell RNA-seq analysis revealed PTN secreted by fibroblasts acting on itself and macrophages via SDC4 ligand in myocardial hypertrophy

**DOI:** 10.1101/2024.06.25.600722

**Authors:** Ke Sheng, Yuqing Ran, Yuting Guan, Pingping Tan, Rongrong Zhang, Songwei Qian, Hongzhou Lin, Huilan Wu, Yongmiao Peng, Yuqing Huang, Zhiguang Zhao, Guanghui Zhu, Weiping Ji, Xiaoling Guo

## Abstract

**Background:** Hypertrophic cardiomyopathy (HCM) is characterized by massive myocardial hypertrophy, which is the most frequent cause of sudden death and can lead to heart failure (HF) or stroke. The objective of this study was to explore the communication network among various cells in the heart of pathological HCM derived from transverse aortic constriction (TAC) mouse model, and investigate the potential mechanism through data mining, biological informatics analysis, and experimental validation.

**Methods and Results:** The integrated analyses including CellChat, Seurat, gene ontology (GO), pseudo-time trajectory analysis, and weighted gene co-expression network analysis (WGCNA) were performed based on the single-cell RNA-seq data (scRNA-seq). I*n vitro* tests were conducted to verify bioinformatic analysis findings through enzyme-linked immunosorbent assay (ELISA), real-time quantitative PCR (RT-qPCR), Edu staining, and transwell assay. *In vivo* tests were also performed to further verify bioinformatic analysis findings by western blot and immunofluorescence assays based on our established TAC mouse model with myocardial hypertrophy. Our results showed that in the heart of TAC mouse, the interaction between cardiac fibroblasts and macrophages was most common, and the increasing pleiotrophin (PTN) secreted by cardiac fibroblasts could promote themselves proliferation or invasion as well as stimulate macrophage activation to release inflammatory cytokines, such as TNF-α, IL-6, Cox-2, Cd83, Egr2, and IL-10 through acting on its ligand recombinant Syndecan 4 (SDC4), which may affect cardiomyocyte normal function and eventually cause HCM. This study first demonstrated that PTN derived from cardiac fibroblasts may act on SDC4 to play crucial role in myocardial hypertrophy, which may be a potential therapeutic targets for patients with pathological HCM.

**Conclusions:** In this study, the complex interaction network between cardiac fibroblasts and macrophages of TAC mice based on the scRNA-seq data was investigated, and we found that the increasing PTN secreted by cardiac fibroblasts under cardiac pressure overload could promote themselves proliferation or invasion as well as stimulate macrophage activation to release inflammatory cytokines through acting on SDC4 ligand, which may affect cardiomyocyte normal function and eventually cause HCM. In addition, our study suggested that PTN derived from cardiac fibroblasts may act on SDC4 ligand to play crucial role in myocardial hypertrophy, which may be a potential therapeutic targets for patients with pathological HCM.

## Introduction

Hypertrophic cardiomyopathy (HCM) is characterized by massive myocardial hypertrophy and dynamic left ventricular outflow obstruction, which is the most frequent cause of sudden death in young people and can lead to heart failure (HF) or stroke^[1]^. HF is a common outcome of multiple cardiovascular diseases, and has a poor prognosis^[2]^. Despite advances in HF treatment, the incidence is still increasing^[3]^. In response to hemodynamic factors such as mechanical stress or pressure overload, cardiomyocytes develop hypertrophy to adapt. When the pressure overload persists, cardiomyocytes undergo maladaptive changes and gradually develop into cardiac fibrosis, ultimately leading to HCM^[4]^. However, the pathological process and underlying mechanism of sustained pressure overload-induced excessive myocardia hypertrophy developing into irreversible fibrosis remain incompletely understood.

There are two forms of myocardial hypertrophy including physiological hypertrophy induced by exercise and pathological hypertrophy induced by abnormal stresses such as hypertension, pressure overload, endocrine disorders, myocardial infarction, and contractile dysfunction caused by genetic mutations of myocardium sarcomere or cytoskeleton^5,6^. Pathological myocardial hypertrophy is accompanied by increased oxygen demand of cardiomyocytes, decreased coronary perfusion pressure, and increased external pressure on coronary microvessels. The increased perfusion requirement is coupled with increased coronary resistance, leading to an imbalance between myocardial oxygen supply and demand, and resulting in hypertrophy relative ischemia/hypoxia^[7]^. In addition, pathological hypertrophy is also associated with decreased capillary density^[8]^. During the adaptation phase of myocardial hypertrophy in mice, hypertrophic stimulation could induce the expression of vascular endothelial growth factor (VEGF) or angiopoietin-2, and blocking VEGF signaling pathway could lead to a decrease in capillary density and early HF^[9]^. Hypertrophy of VEGF deficient mice induced by pressure overload could accelerate the transition from compensatory hypertrophy to failure^[10]^, which suggested that coronary angiogenesis occurs during physiological hypertrophy, and vascular sparsity in pathological hypertrophy may lead to myocardial hypoxia and systolic dysfunction. So far, the classic TAC mouse model can accurately simulate pathological myocardial hypertrophy processes and characteristics of HCM^[11]^.

The heart is composed of different cell types, of which endothelial cells, cardiomyocytes, and cardiac fibroblasts are the most abundant. Communication between these cell types, also known as paracrine signaling, is essential for normal cardiac function but is also crucial in cardiac remodeling and HF^[12]^, and most cells, including the major cell types in the heart, have many ligand-receptor pairs, suggesting that autocrine signaling is a universal phenomenon^[13]^. Autocrine signaling in cardiac remodeling even HF is involved in verious pathophysiological phenomena, such as hypertrophy, fibrosis, angiogenesis, cell survival, inflammation, and so on^[14]^. Myocardial fibroblasts may be activated in some way under stress, transitioning from dormant cells into proliferating cells and then into secreting cells.

In this study, we would explore the network of paracrine and autocrine signals among different cell types in the pathological HCM heart of transverse aortic constriction (TAC) mouse model, and try to investigate the potential mechanism of pathological HCM through biological informatics analysis of single-cell RNA-seq data and *in vitro* and *in vivo* experimental validation.

## Material and Methods

### Study approval

All animal studies and numbers of animals used conform to the Directive 2010/63/EU of the European Parliament and have been approved by the appropriate local authorities (Wenzhou Medical University’s Animal Care and Use Committee).The male C57BL/6 mice were provided by Laboratory Animal Center, Wenzhou Medical University, Wenzhou, China. In all experiments, animals were killed by cervical dislocation under isoflurane anaesthesia (induced by isoflurane inhalation 4.0%).

### Single-cell RNA-seq (scRNA-seq) data acquisition

The scRNA-seq data of GSE137167 was downloaded from the Gene Expression Omnibus (GEO) database (https://www.ncbi.nlm.nih.gov/geo), including CD45^+^ cell samples from the hearts of healthy (sham) and TAC mice (6 per group), and the single-cell mRNA atlas was studied using 10× Genomics technology^[15]^.

### scRNA-seq analysis

The scRNA-seq data for each sample was analyzed separately. Standard processing steps including filtration, highly variable gene identification, dimensionality reduction, and cell clustering were performed using the “Seurat” package in version 4.2.0 R software^[16]^. Subsequently, low-quality cells expressing with less than 300 genes or more than 2500 genes as well as genes expressing with less than 3 cells were excluded to eliminate empty droplets or potential doublets and multiplets. Then, cells with mitochondrial RNA percentage >10% were considered poor quality and filtered. Cells with hemoglobin RNA percentage >0.1% were considered red blood cells and removed. To normalize the data, gene expression levels of the remaining cells were multiplied by 10,000 and logarithmically converted. Highly variable genes were selected using standard deviation and used for downstream analysis. To scale the data, the ScaleData function in the Seurat package set the average expression of genes between cells to 0 and their variance to 1. Principal component analysis (PCA) was performed on the top 3,000 highly variable genes (HVGs), and 13 principal HVGs were selected. The cells are divided into different clusters using Louvain’s algorithm. The *t*-distributed stochastic neighbor embedding (*t*SNE) was used for two-dimensional visualization of clustered cells. The FindAllMarkers function was used to test and identify differentially expressed genes (DEGs) within each cluster. |Log2FC|> 0.5 and adjusted *p* value (pvals_adj) < 0.05 were considered significant DEGs. The “singleR” package was used to annotate cells^[17]^. Clusters annotated as fibroblasts were selected, and cardiac fibroblasts from sham and TAC mice were merged using Harmony package to minimize batch effects^[18]^. The FindMarkers function was then used to calculate DEGs of cardiac fibroblasts between sham group and TAC group.

### Intercellular communication analysis

The “CellChat” package in R software was used to analyze and visualize cell-cell communication between different types of cells based on L-R interactions^19^. It analyzes the expression abundance of ligand-receptor interactions between two cell types based on the expression of the receptor in one type of cells and ligand expression in the other. By analyzing the expression abundance of ligand-receptor interactions, we obtain the number of interactions between the two cell types, allowing an initial evaluation of cell-cell communication. In summary, the standardized scRNA-seq data of the “Seurat” package is imported into the “CellChat” package.

### Gene ontology (GO) enrichment analysis

Gene Ontology (GO) is a standardized classification system for gene function, consisting of three ontologies such as Molecular Function (MF), Cellular Component (CC), and Biological Process (BP). To infer the biological functions of SDC4, all genes in the module containing SDC4 were analyzed using GO enrichment analysis in the clusterProfiler package^[20]^, which calculates the number of genes for each term. The GO terms of the module genes were defined by hypergeometric tests.

### Single-cell pseudo-time analysis

Monocle 3 was used for pseudo-time trajectory analysis of single cells. The differentiation pathways between each macrophage subtype and fibroblast as well as changes in target gene expression along pseudo-time were predicted, without prior knowing the differentiation timing or direction. The analysis workflow was executed based on the official guidelines of Monocle 3 (https://cole-trapnell-lab.github.io/monocle3).

### Weighted gene co-expression network analysis (WGCNA)

Weighted gene co-expression network analysis (WGCNA) in the R package^[21]^ was used to identify co-expressed gene modules and explore the relationship between gene networks and phenotype of interest. The analysis followed these criteria, such as soft threshold of 3, minimum module size of 50, and cutting height of 0.25 (MEDissThres = 0.25). Genes with similar expression patterns are combined into a single module. Finally, we identified the module where *Sdc4* was located for GO enrichment analysis to predict its function.

### Cell culture

The mouse macrophage cell line RAW264.7 (ATCC, VA, USA) was cultured in dulbecco’s modified eagle medium (DMEM) with 10% fetal bovine serum (FBS, Gibco, CA, USA) at a 37°C and 5% CO_2_ incubator. The mouse cardiac fibroblast cell line L929 (ATCC, VA, USA) were cultured in RPMI 1640 medium containing 10% FBS and 1% penicillin/streptomycin (P/S, Gibco, CA, USA) at a 37°C, 5% CO_2_ incubator or at a 37°C, 5% CO_2_ hypoxic incubator with 3% O_2_.

### Detection of cell proliferation viability and invasion ability

CCK-8 assay was used to detect cell viability in different groups. Briefly, cells were seeded in a 96-well plate at a density of 1×10^4^ cells/well, and cultured at 37°C, 5% CO_2_ incubator for 24 h with or without PTN treatment. Then, 10 μL of CCK-8 solution diluted 10 times by culture medium was added into each well and incubated for 1.5 h at 37°C incubator. Finally, the absorbance at 450 nm was measured using a microplate reader. In addition, EdU staining using BeyoClickTM EdU-488 cell proliferation assay kit (C0071S, Beyotime, Beijing, China) was performed to further assess cell proliferation activity in different groups. EdU positive cells with green fluorescence were proliferative cells.

Transwell assay was conducted to assess cell invasion and migration ability in different groups. Briefly, the transwell chambers were placed in a 24-well plate. 3×10^4^ cells were seeded in the upper compartments of chambers containing 200 μL of serum-free medium supplemented with 1% BSA to maintain osmotic pressure. The lower compartments of chambers were filled with 400 μL of complete medium supplemented with or without 0.1 μg/mL recombinant mouse PTN^[22]^. After incubation for 24 h, cells were scraped with cotton swabs, then fixed with 4% paraformaldehyde, and finally stained with 0.1% crystal violet. Images were captured by an inverted optical microscope (OLYPAS, Japan).

### Quantitative PCR (qPCR) analysis

Total RNAs were extracted from different group cells using Trizol reagent (Invitrogen, CA, USA), and then were reverse-transcribed into cDNAs using cDNA Synthesis SuperMix (Vazyme, Nanjing, China). Subsequently, these cDNAs were used as templates for RT-qPCR with SYBR Green SuperMix (Vazyme, Nanjing, China). The reaction mixture containing 10 μL SYB Green Mix, 0.5 μL forward primer, 0.5 μL reverse primer, 1 μg diluted cDNA, and 5-8 μg ddH_2_O. The reaction process was 95 ℃ for 3 min, followed by 40 cycles of 95°C for 20s, 60°C for 30s. The relative expression of genes was normalized against *Gapdh*. The quantification was performed using the comparative Ct (2^-ΔΔCt^) method. The primer sequences are listed in Table 1.

### Enzyme-linked immunosorbent assay (ELISA)

In order to detect the level of PTN secreted by cardiac fibroblasts cultured at the hypoxic environment for 0 h, 24 h, 48 h, and 72 h to simulate the cardiac pressure overload, enzyme-linked immunosorbent assay (ELISA) kit was used to detect the PTN levels of supernatant extracted at different time points according to the manufacturer’s instructions (Chemicon CA, USA). Briefly, 50 μL of assay diluent was added into pre-coated wells. The plates were incubated for 2 h at room temperature, and then were washed 5 times with washing buffer. 100 μL of peroxidase-conjugated IgG anti-PTN solution was added into each well. Then, the plates were washed with washing buffer 5 times again. Then, 100 μL of substrate buffer was added into each well and kept in dark for 30 min at room temperature. The reaction was stopped using 50 μL of stop solution. The absorbance is measured at 450 nm wavelength (OD value), and the concentration of PTN is calculated using a standard curve.

### Transverse aortic constriction (TAC) surgery

Male C57BL/6 mice weighing 20-25 g were housed in a Specific Pathogen-Free (SPF) environment, and maintained in a 12 h dark/light cycle at the temperature of 23 ± 2°C and relative humidity of 45% to 55%. Water and food were accessed ad libitum.Transverse aortic constriction (TAC) surgery was performed in a sterile operating room, and the surgical site was disinfected using 75% alcohol. Mice were anesthetized in a chamber with 2% isoflurane mixed with 100% oxygen at a flow rate of 0.5-1.0 L/min. The mouse fur from the neck to the middle of the chest was shaved. Thoracotomy was conducted on the second rib under a surgical microscope, and a rib spreader was used to retract the sternum. Simultaneously, a 6.0 silk suture on a 27G needle was placed between the innominate artery and the left carotid artery, and the transverse aorta was identified and sutured. Two rapid knots were tied, and the needle was quickly withdrawn to create a constriction with a diameter of 0.4 mm. In sham mice, the entire procedure was same, except for the aortic ligation.

### Hematoxylin and Eosin (H&E) staining

The hearts from each group were fixed in a 4% paraformaldehyde solution for 24 h. Subsequently, the dehydrated tissues were embedded in paraffin and cut into slices with a thickness of 5 μm. The myocardial structural changes were evaluated was by Hematoxylin and Eosin (H&E) staining.

### Echocardiography

Mice were subjected to anesthesia with a mixture of 2% isoflurane, and then were treated with 0.5-1.0 L/min of 100% oxygen for 2 min. The fur on the anterior chest was shaved, and an even application of ultrasound coupling agent was made on both the 550-type probe and the chest area. Parameters including IVSd, LVIDd, LVPWd, IVS, LVIDs, LVPWs, EF, and FS were harvested using the VisualSonics high-resolution Vevo 2100 system (FUJIFILM VisualSonics, Tokyo, Japan.

### Immunofluorescence

The paraffin sections of mouse hearts in each group were baked at 65°C for 30 min, dewaxed with xylene, dehydrated by a gradient of alcohol, and repaired with sodium citrate antigen. After permeabilization with Triton X-100, the sections were incubated with primary antibodies (listed in Table 2) at 4°C ovrenight. The sections were rinsed with PBS for three times, then were incubated with AlexFluor 594 AffiniPure goat anti-rabbit IgG or AlexFluor 488 AffiniPure goat anti-mouse IgG at room temperature for 1 h. Finally, DAPI containing anti-fluorescence quencher was used for nuclear staining, Images were collected under an Olympus fluorescence microscope (Tokyo, Japan).

### Western Blot

The left ventricular tissues of mice in each group were homogenized in RIPA lysis buffer containing 1 mM PMSF (Beyotime, Shanghai, China). Protein concentration was determined using the BCA protein assay kit (Beyotime). Protein samples were isolated using 10% sodium dodecyl sulfate-polyacrylamide (SDS) gel, and then transferred to polyvinylidene difluoride membranes (Millipore, MA, USA). After blocking with 5% nonfat milk (Beyotime) for 2 h, the membranes were incubated overnight at 4°C with primary antibodies (listed in Table 2). After washing three times with TBST, the membranes were incubated at room temperature for 1 h with secondary antibodies (goat anti-rabbit or goat anti-mouse) conjugated with horseradish peroxidase. The protein bands were detected using the ECL detection kit (Beyotime).

### Statistic analysis

The data is presented as mean ± standard deviation (SD). All statistical analyses were conducted using the statistical package in GraphPad Prism 9 (version 6.02). Student’s *t*-test was used to compare two experimental groups with unpaired data. A *p*-value <0.05 was considered statistically significant.

## Results

### Overview of different cell types in CD45+ cell samples from TAC mice hearts

The processed scRNA-seq data were screened using standard processing workflow. We define mitochondria as cells in which the expression of mitochondrial genes accounts for more than 10% of the total gene expression. For the identification of red blood cells, we examined genes such as *Hba-a1*, *Hba-a2*, *Hbb*, *Hbb-bh2*, *Hbb-b2*, and *Hba*. Cells with a proportion of these genes exceeding 0.1% of the total gene expression are classified as red blood cells. Cells with nFeature_RNA less than 300 are considered potential empty droplets or dead cells. Simultaneously, we exclude cells with nFeature_RNA greater than 7500 or nCount_RNA greater than 10000, as these cells may indicate the presence of two or more cells within a droplet (Fig. S1-S4). The single-cell data of TAC group was clustered based on 3000 HVGs (Fig. 1B), resulting in 25 clusters (Fig. 1A). Next, these clusters were annotated using the single R package, and their cell types were determined based on marker genes (Fig. 1D). Finally, it was determined that the cell types in CD45^+^ cell samples from TAC mouse hearts included NK cells, macrophages, monocytes, fibroblasts, granulocytes, and endothelial cells (Fig. 1C). Based on our analysis, we have identified the cell types and corresponding clusters of cells in TAC group, which will facilitate further analysis.

**Fig. 1.**
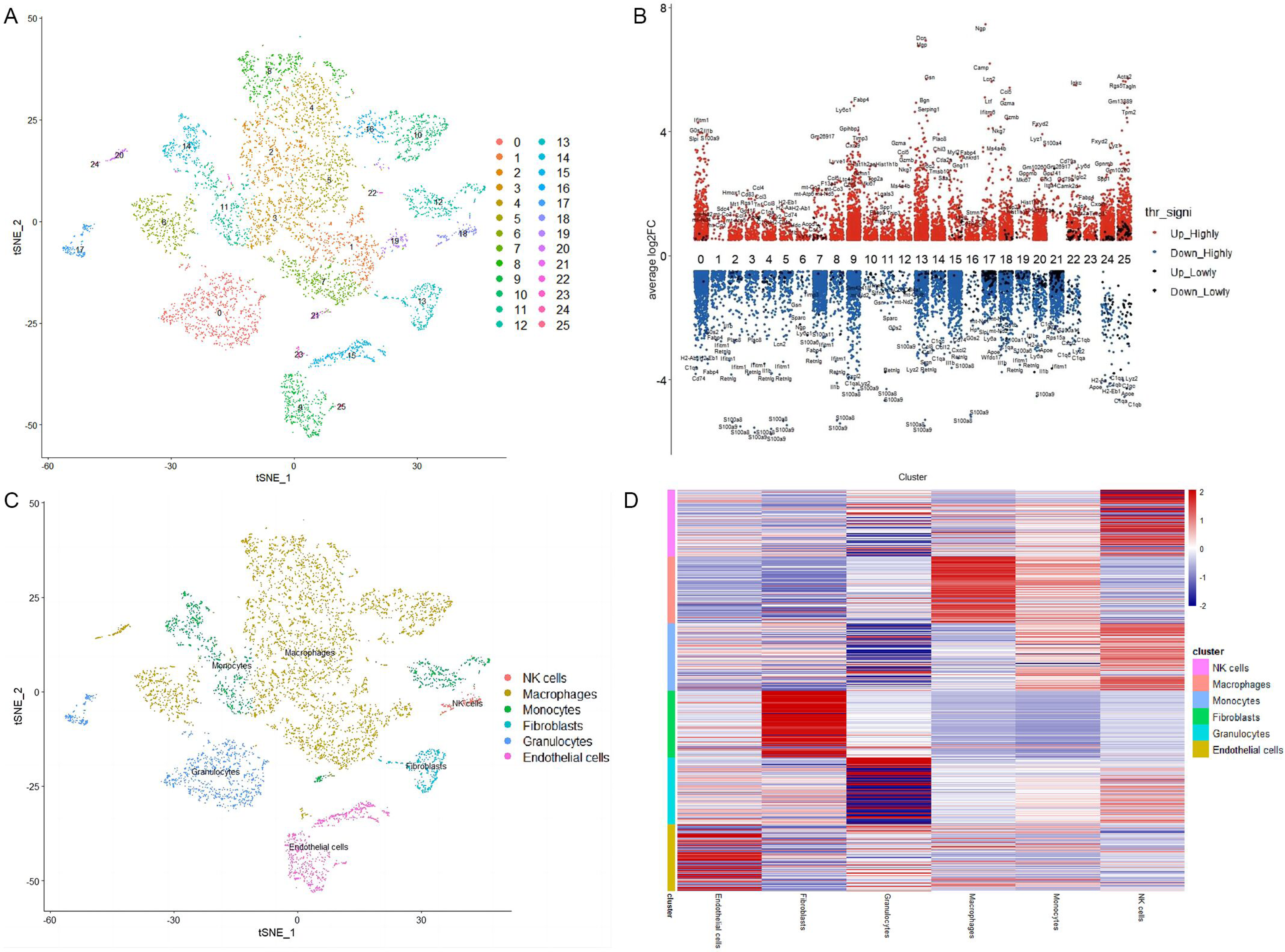
Clustering and annotation of single-cell sequencing data. (A) This visualization depicts the t-SNE plot, which illustrates distinct clusters of cells derived from scRNA-seq data. Each unique cell cluster is represented by a different color. (B) For each cell cluster population, the differential gene expression is depicted, with blue indicating downregulation and red indicating upregulation. (C) According to the marker genes, the corresponding cell types for each cell cluster are determined, and each color represents a specific cell type. (D) Differential expression heatmaps of marker genes for each cell type, where log FC > 0 is represented in red and log FC < 0 is represented in blue.

### The cell-cell communication analysis

Based on the expression levels of ligand/receptor pairs between these six types of cells, the network of cell-cell communication was constructed using CellChat software (Fig. 2A). A total of 403 ligand-receptor pairs have been identified (Supplement data 1). The interaction number, which was indicated by line weights in network planning or shade of color in heatmap, was plotted in terms of ligand-receptor pairs, respectively (Fig. 2B). The results showed that mouse cardiac fibroblasts had the most interactions with themselves or monocytes/macrophages. Next, we need to investigate the specific ligand-receptor pairs between fibroblasts and themselves as well as between cardiac fibroblasts and macrophages. Then, some types of ligands from these cells were displayed. Cardiac fibroblasts had all ligands included in pattern 1 (Fig. 2C), of which PTN mainly acted on itself and followed by monocytes/macrophages (Fig. 2D). We believe that the PTN pathway is representative, and it may have certain biological effects on both fibroblasts and macrophages.

**Fig. 2.**
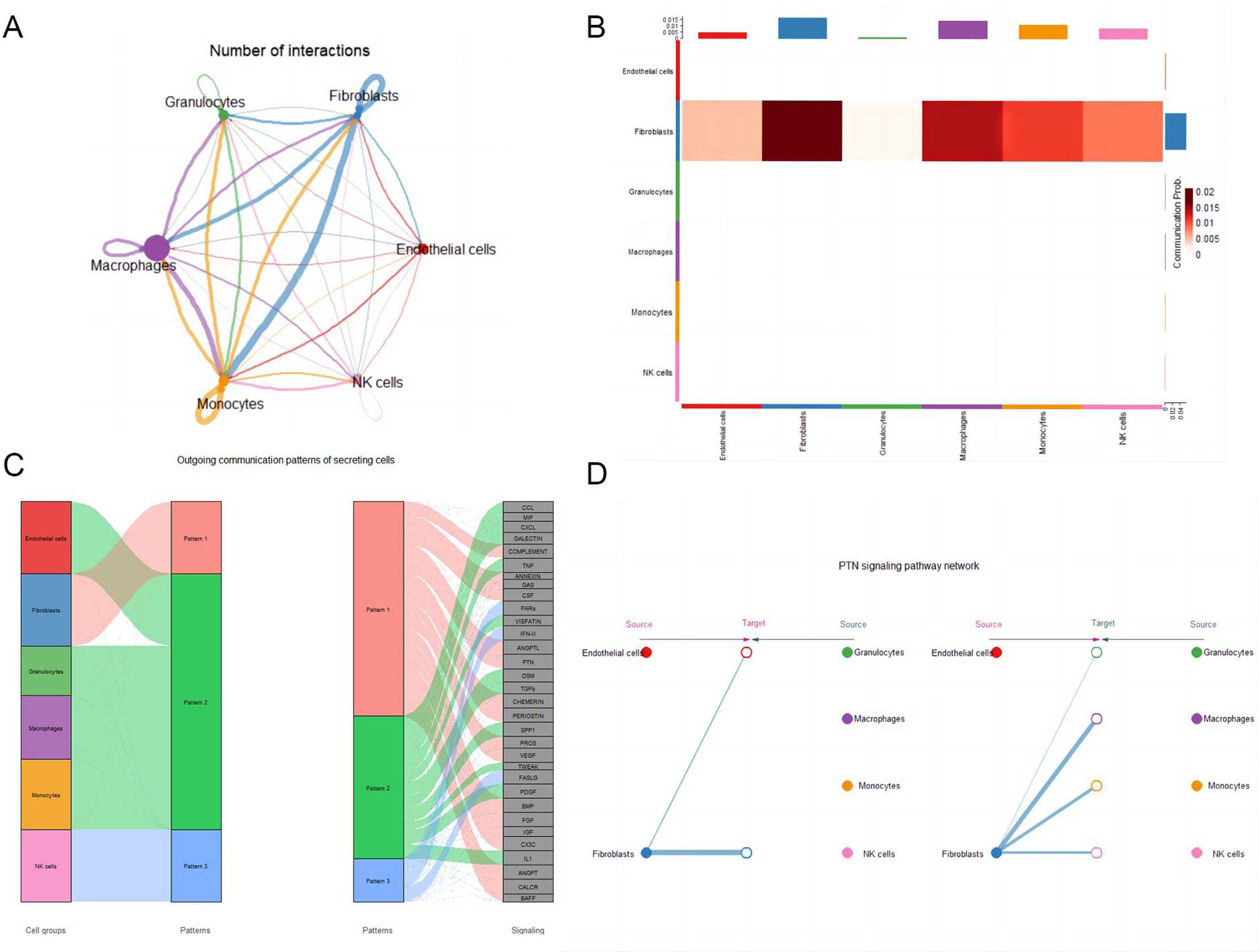
The interaction between different types of cells. (A) The communication network among various types of cells. The thicker the connections between cells, the more numerous the ligand-receptor pairs. Conversely, the thinner the connections, the fewer the ligand-receptor pairs. (B) The heatmap of ligand-receptor pairs between different cell types, with the horizontal and vertical axes representing specific cell types. (C) The left panel represents a collective term for the total ligands secreted by each cell type, referred to as a pattern. The right panel displays the specific ligands secreted by each cell type, grouped together as patterns, revealing which ligands are secreted by different cell types. (D) The interaction strength of the PTN pathway between different cell types. The thickness of the lines represents the number of ligand-receptor pairs involved in the PTN pathway. Different cell types are represented by circles of different colors, with solid circles indicating the source and hollow circles indicating the target.

### Definition of interactions for ligand/receptor pairs in the PTN pathway and evaluation of PTN secretion by fibroblasts under hypoxia conditions

The ligand/receptor pairs of the PTN pathway included PTN-NCL, PTN-SDC2, PTN-SDC3, and PTN-SDC4 (Fig. 3A). The gene expression levels of these members including *Ptn*, *Sdc2*, *Sdc3*, *Sdc4*, and *Ncl* in mouse different cardiac cell types such as endothelial cells, cardiac fibroblasts, granulocytes, macrophages, and monocytes were analyzed. Interestingly, PTN as a ligand is only expressed in mouse cardiac fibroblasts.

**Fig. 3.**
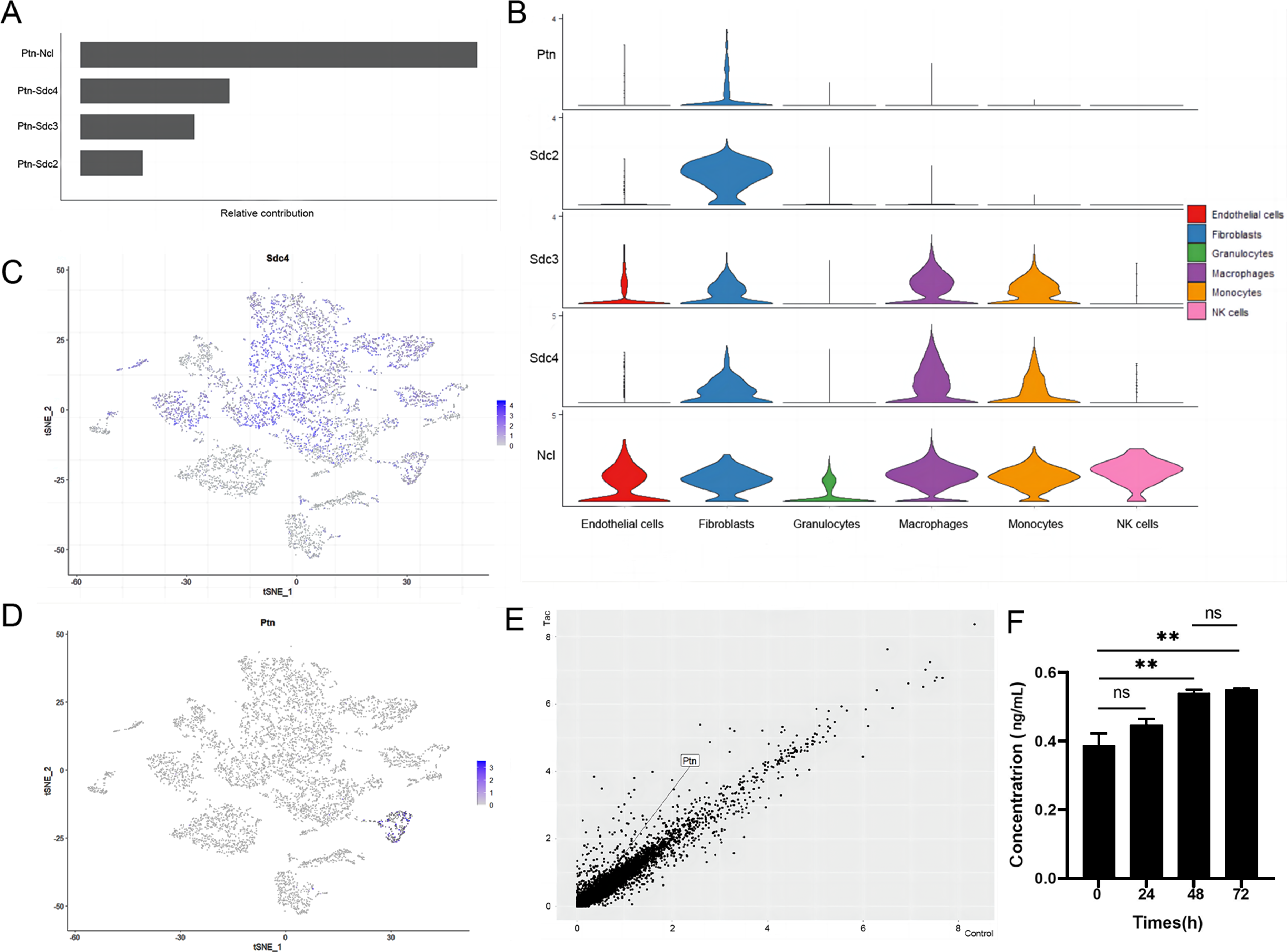
The PTN pathway between fibroblasts and other types of cells. (A) Known ligand-receptor pairs and their weights in the PTN pathway. (B) Violin plot of the expression levels of ligand and receptor members in various cell types in the PTN pathway. (C) Expression level distribution of *Sdc4* across all cell clusters. (D) Expression level distribution of *Ptn* across all cell clusters. (E) Volcano plot illustrating the differential gene expression between TAC (treatment) fibroblasts and control fibroblasts. (F) ELISA measurement of the secretion level changes of PTN in vitro fibroblasts with prolonged hypoxia exposure time(n=3). Mean±SD,*p<0.05,**p<0.01,***p<0.0001 indicate significant differences,and ns>0.05 means no significant difference.

As receptor of PTN, SDC2, SDC3, SDC4, and NCL are abundantly expressed in cardiac fibroblasts or monocytes/macrophages (Fig. 3B). In terms of gene expression distribution, we found that *Sdc4* is widely expressed and highly present in various macrophage clusters, but it is also present in single cardiac fibroblast cluster (Fig. 3C), and *Ptn* is also mainly distributed in cardiac fibroblast cluster (Fig. 3D). Moreover, among the four receptors in the PTN pathway, only SDC4 is expressed in cardiac fibroblasts, monocytes, and macrophages. In addition, we integrated the expression matrix of control and TAC groups and used the harmony package to remove batch effects to calculate the different expression levels of *Ptn* between sham and TAC fibroblasts. The results showed that compared with sham group, the level of *Ptn* was up-regulated in TAC group, with a logFC of 0.46 (Fig. 3E). To validate this result, PTN levels in the supernatant of mouse cardiac fibroblasts cultured at hypoxia condition for 0 h, 24 h, 48 h, and 72 h were detected by ELISA. The results showed that with the extension of hypoxia time, the level of PTN in the supernatant increased and reached the plateau at 48 h or 72 h, which was significantly different from that at 0 h (Fig. 3F). Therefore, we believe that the pathway involving PTN secretion by cardiac fibroblasts and its action on both self and macrophage SDC4 is unique. The results of bioinformatics analysis and *in vitro* cell experiments consistently demonstrate an increase in PTN expression, suggesting that the PTN pathway may play a role in the process of pressure overload.

Meanwhile, the level of *Sdc4* in macrophages of TAC group was upregulated compared with sham group (Supplement data 2). Monocle 3 was used to calculate the developmental trajectories of cardiac fibroblasts in TAC group (Fig. S5), and the expression changes of *Ptn* over simulated time were displayed based on their developmental states (Fig. S6). The expression level of *Ptn* in cardiac fibroblasts increased slowly with simulation time, which was consistent with ELISA result, suggesting that PTN content may exhibit a similar trend with disease progression.

### Prediction of Sdc4 function using WGCNA approach and validation by qPCR

The expression matrix of control group and TAC group was integrated, and the batch effects was removed using Harmony package. WGCNA analysis (Fig. 4A) was performed on the macrophages in sham and TAC groups to obtain phenotypic related modules (Fig. 4B). These modules are essential collections of genes that are co-expressed. *Sdc4* was included in the module with the strongest phenotype correlation (Supplement data 3). To evaluate the biological function of *Sdc4* and its potential effects on macrophages, we intersected all genes in this module with DEGs in sham and TAC macrophages (Fig. 4C), and then performed GO molecular function enrichment analysis (Fig. 4D). The results showed that the functions of intersecting genes were mainly enriched in the positive regulation of response to external stimuli, cytokine-mediated signaling pathway, and bacterial-derived molecules, suggesting that *Sdc4* may be involved in promoting the inflammatory response of macrophages.

**Fig. 4.**
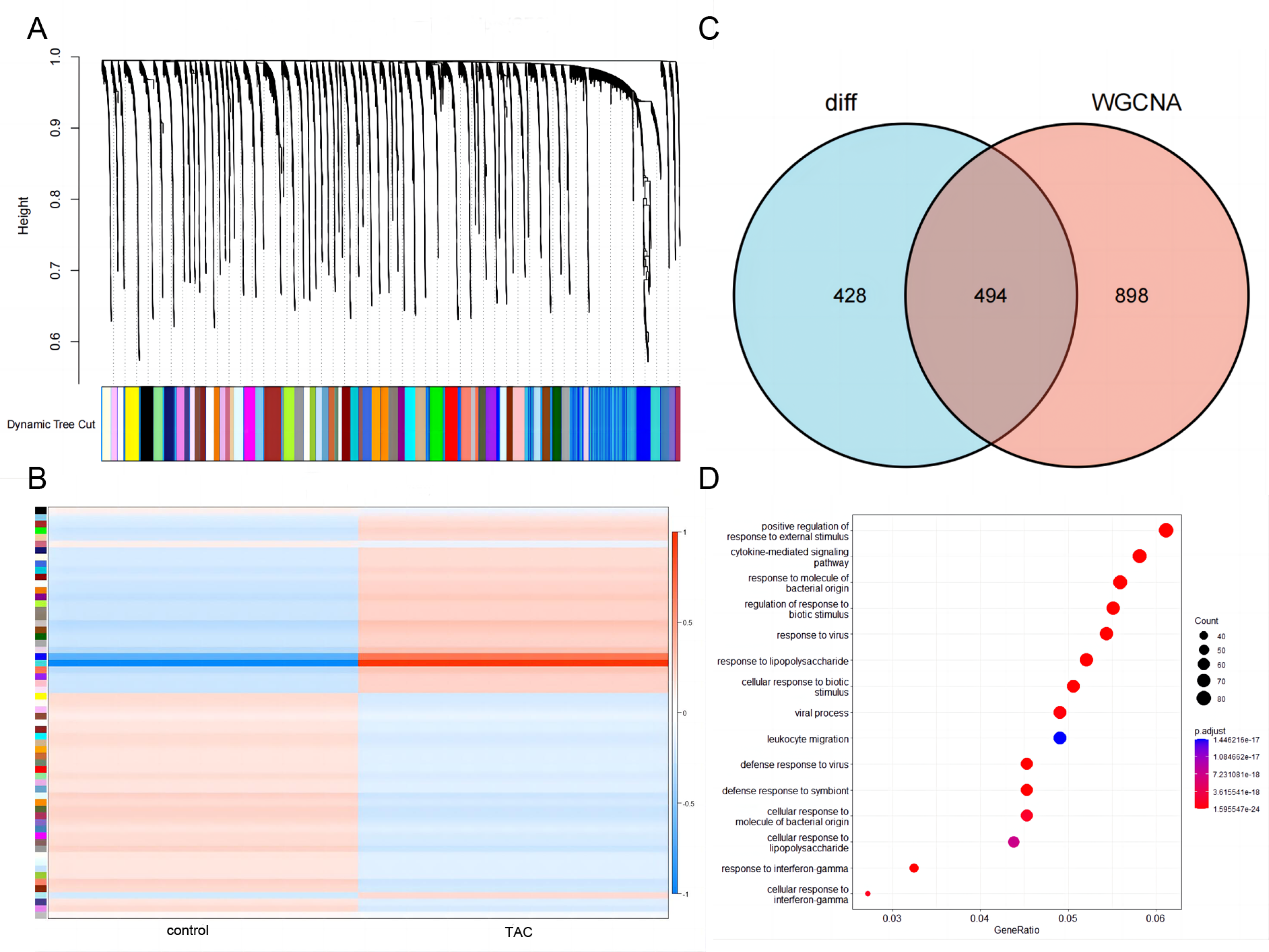
Functional prediction of Sdc4 in macrophages. (A) Clustered dendrogram of all differentially expressed genes based on dissimilarity measurement (1-TOM). The color band represents the outcomes obtained through automated single-block analysis. (B) Heatmap of the correlation between the module eigengenes and clinical traits of TAC. (C) Venn diagram depicting the intersection of differentially expressed genes (DEGs) between the TAC group and the sham group in macrophages, as well as the genes included in the module containing *Sdc4*. (D) Bubble plot illustrating the Gene Ontology (GO) enrichment results of the intersection genes in the Venn diagram in terms of their molecular functions.

To validate this hypothesis, mouse macrophages (RAW264.7) were incubated in complete culture medium containing 0.1 μg/mL recombinant mouse PTN for 24 h. We then selected several inflammatory markers such as *Cox-2*, *IL-6*, and *TNF-α* (Fig. 5A-5C) as well as some markers that partially characterize macrophage polarization toward M2 phenotype such as *EGR-2*, *IL-10*, and *CD83* (Fig. 5D-5F) to perform qPCR to detect their mRNA expression levels. The results showed that compared with control group (0 μg/mL PTN), these inflammatory markers were elevated and these markers of M2 phenotype macrophages were downregulated. The qPCR results confirmed our hypothesis that the function of PTN in macrophages was consistent with the bioinformatics predictions.

**Fig. 5.**
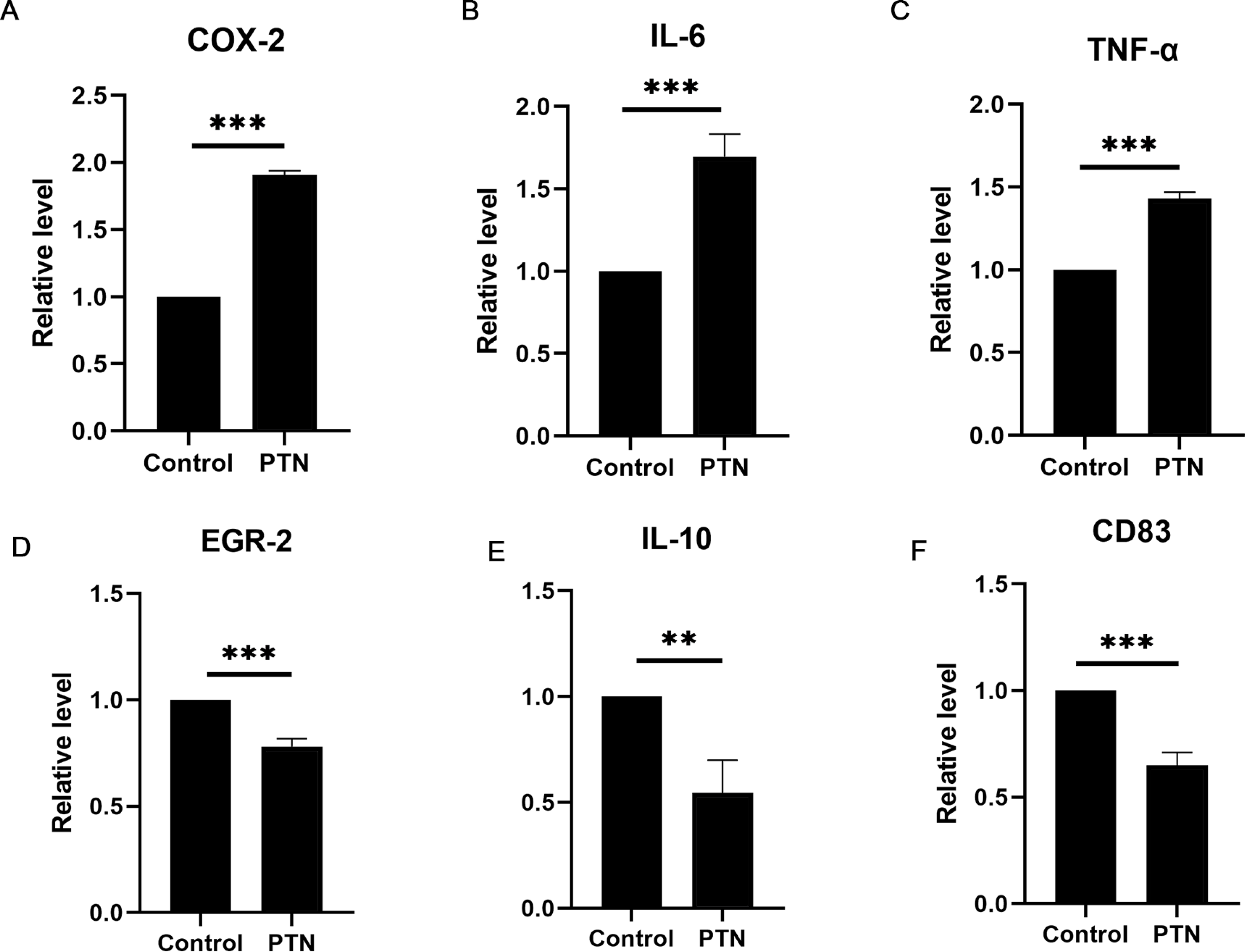
Bar chart depicting the qPCR results of markers for M1 macrophages and M2 macrophages. (A) Relative expression levels of *cox2* mRNA in the PTN group and control group(n=3). (B) Relative expression levels of *Il6* mRNA in the PTN group and control group(n=3). (C) Relative expression levels of *Tnfa* mRNA in the PTN group and control group(n=3). (D) Comparative mRNA expression levels of *Egr2* between the PTN group and the control group(n=3). (E) Comparative mRNA expression levels of *Il10* between the PTN group and the control group(n=3). (F) Comparative mRNA expression levels of *Cd83* between the PTN group and the control group(n=3). Mean±SD,*p<0.05,**p<0.01,***p<0.0001 indicate significant differences,and ns>0.05 means no significant difference.

### PTN enhances the proliferation viability and invasion ability of fibroblasts

PTN is secreted mainly by cardiac fibroblasts, and both macrophages and cardiac fibroblasts themselves can express PTN ligand. It is well known that under pressure overload, cardiac fibroblasts in the heart undergo a transition from dormant cells to proliferating cells and then to secreting cells^23^. To investigate the potential effects of PTN on cardiac fibroblasts, mouse cardiac fibroblasts were cultured in complete culture medium containing 0.1 μg/mL recombinant mouse PTN for 24 h. Their proliferative activity was examined using CCK8 assay and EdU staining (Fig. 6A), and their invasive ability was assessed by Transwell assay (Fig. 6C). The results showed that compared with control group (0 μg/mL PTN), PTN could markedly increase the numbers of positive EdU staining cells with green fluorescence (Fig. 6B), and significantly promote the cell viability (Fig. 6E). Moreover, the Transwell assay also showed that compared with control group (0 μg/mL PTN), PTN could dramatically accelerate the invasive ability of fibroblasts (Fig. 6D). Therefore, these data suggested that PTN may be involved in the activation of cardiac fibroblasts in the heart under stress overload.

**Fig. 6.**
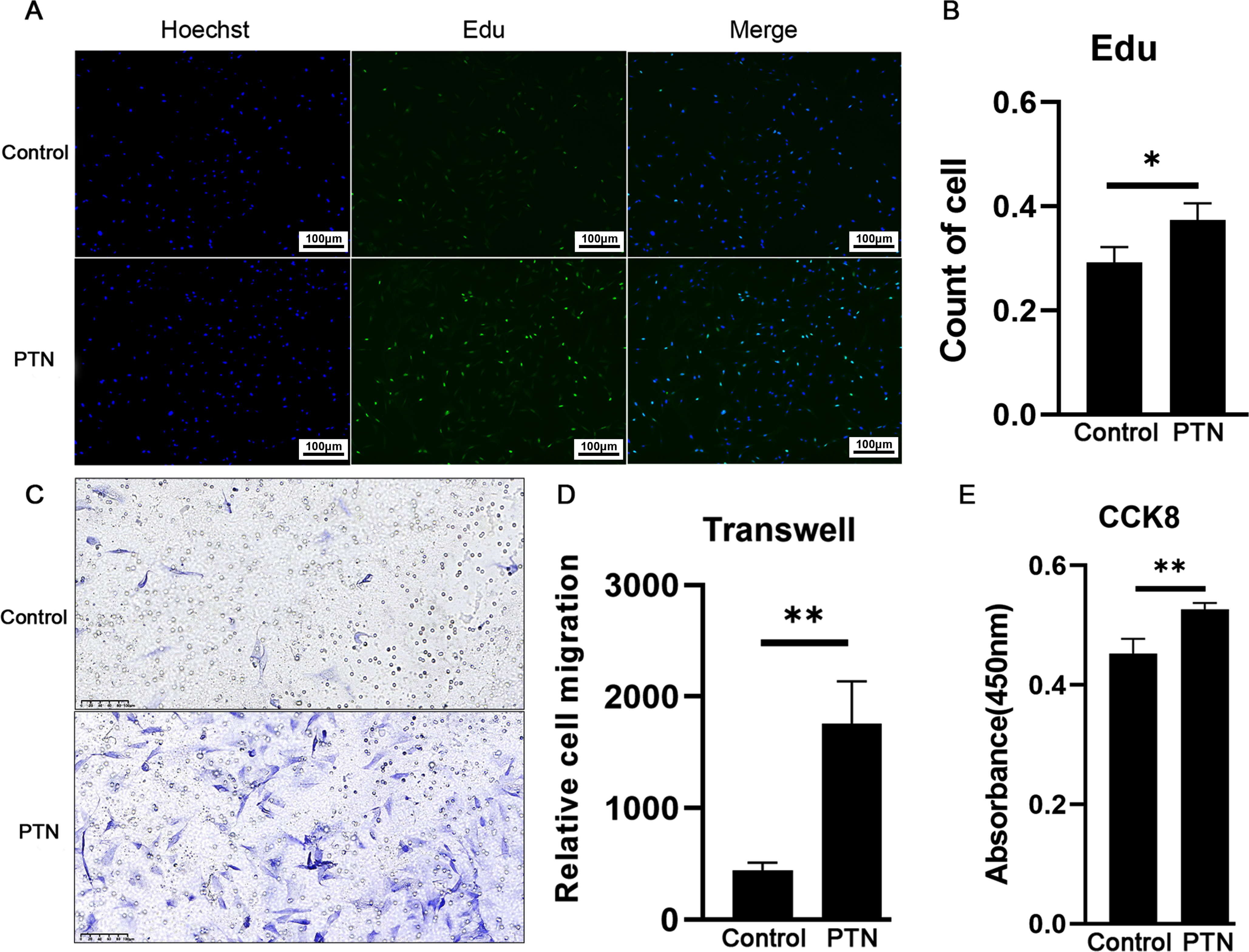
The effect of PTN on proliferation and migration ability of fibroblasts. (A) Fluorescent images of EdU staining in the control group and PTN group(n=3).Scale bar 100μm. (B) Comparison of cell counts in the control group and PTN group after EdU staining. (C) Photographs of transwell crystal violet staining in the control group and PTN group(n=3).Scale bar 100μm. (D) A bar graph illustrating the migration ability of the control group and PTN group cells as assessed by a cell counting assay. (E) A bar graph representing the cell viability of the control group and PTN group measured by CCK-8 assay. Mean±SD,*p<0.05,**p<0.01,***p<0.0001 indicate significant differences,and ns>0.05 means no significant difference.

### PTN is significantly co-localized with fibroblasts in mouse hearts

Transverse aortic constriction (TAC) surgery was performed to induce myocardial hypertrophy in mice. Echocardiography was conducted at 2 and 4 weeks post-operation to assess the cardiac ultrasound indexes in each group (Fig. 7A). The results showed that the cardiac ultrasound indexes including IVsd, LVPWd, IVSs, LVPWs, EF, and FS in TAC group at 2 or 4 weeks after surgery were significantly different from these of sham group (Fig. 7B). At 4 weeks post-operation, the size of the mouse hearts in TAC group was significantly larger than that of sham group (Fig. 7D). HE staining results showed that the size of cardiomyocytes in TAC group increased significantly compared with sham group (Fig. 7C). Additionally, heart index (heart weight/body weight) and interventricular septum thickness were significantly higher in TAC group compared with sham group (Fig. 7E). These data suggested that TAC mouse model has been successfully established for follow-up experimental verification *in vivo*.

**Fig. 7.**
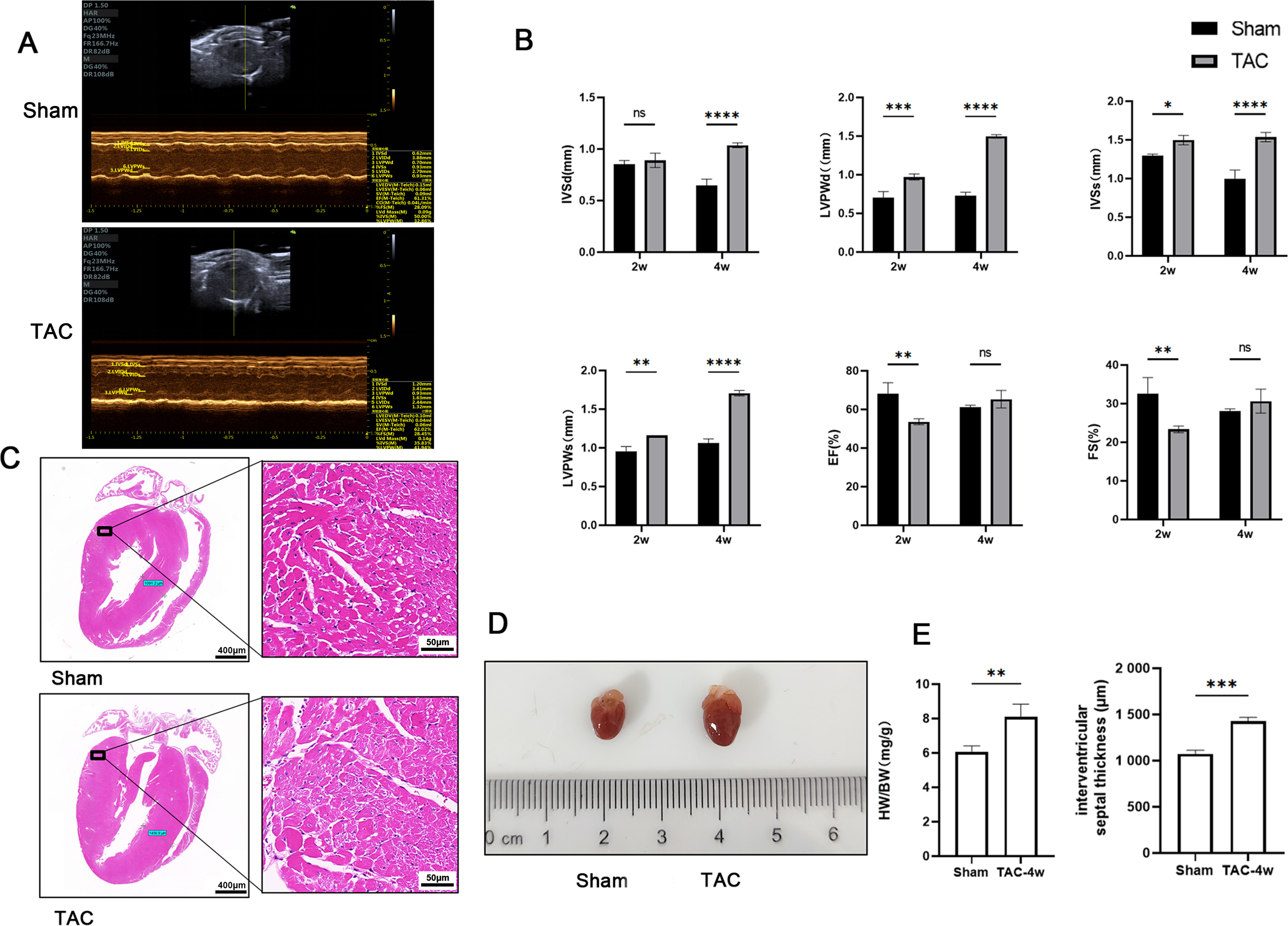
Cardiac function and histopathological evaluation after TAC modeling. (A) The ultrasound images of mice in TAC(n=3) group and control group at two weeks. (B) The data of IVsd, LVPWd, IVSs, LVPWs, EF, and FS in TAC group and sham group at two and four weeks. (C) HE staining of heart sections in TAC group and sham group at four weeks post-modeling.Scale bar 400 μm and 50 μm respectively. (D) The hearts of mice in TAC group and sham at four weeks post-modeling. (E) Comparison of heart index and interventricular septum thickness comparison between TAC group and sham group.Mean±SD,*p<0.05,**p<0.01,***p<0.0001 indicate significant differences,and ns>0.05 means no significant difference.

The mouse heart paraffin sections of TAC and sham group were subjected to immunofluorescence double staining for vimentin (a fibroblast marker) and PTN. There was a significant co-localization of these two proteins in mouse heart (Fig. 8A). Statistical analysis of fluorescence images showed that the fluorescence intensity of both vimentin and PTN in TAC group were higher than that of sham group, and the fluorescence intensity ratio of PTN/DAPI was significantly higher in TAC group compared with sham group, reflecting an overall elevation in PTN expression in the TAC group. Moreover, the fluorescence intensity ratio of PTN/vimentin was also significantly higher in TAC group compared with sham group, indicating an increased expression of PTN in fibroblasts in TAC group (Fig. 8B). Western blot was performed to detect PTN expressions in heart tissues from TAC group and sham group (Fig. 8C), and the result showed a significantly higher total PTN in TAC group than that of sham group (Fig. 8D). These findings suggested that the cardiac fibroblasts could secrete more PTN induced by TAC surgery, which may activate macrophages to release inflammatory cytokines through acting on its ligand SDC4 to affect cardiomyocyte normal function.

**Fig. 8.**
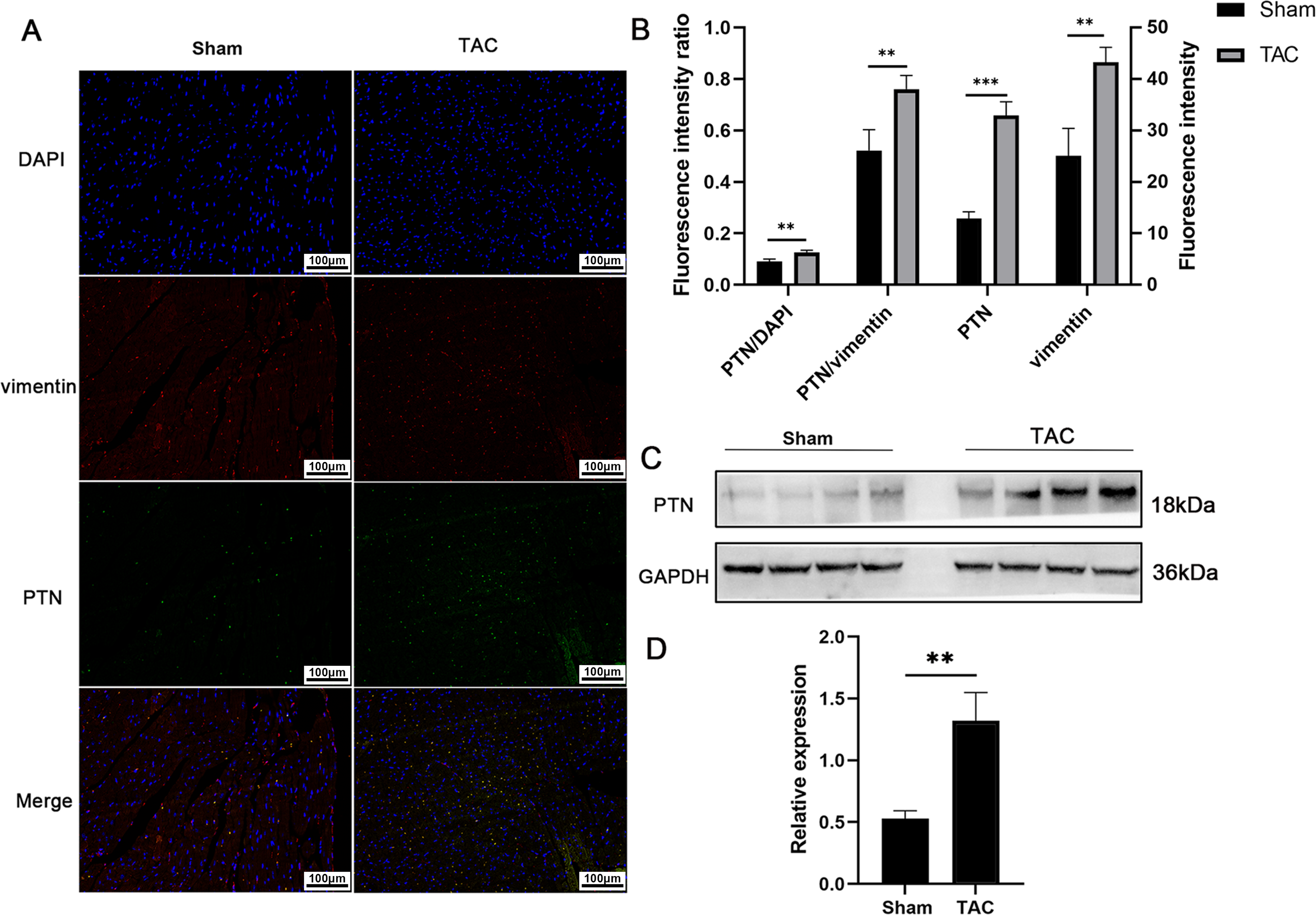
The cardiac tissues of mice exhibit co-localization of PTN and vimentin. (A) Immunofluorescence double labeling of PTN and vimentin in cardiac sections of mice from TAC group and sham group(n=3).Scale bar 100 μm. (B) The quantitative fluorescence intensity of PTN and vimentin in TAC group and sham group. (C) The detection of PTN expression by western blot in TAC group and sham group(n=3). (D) The quantification of PNT bands in TAC group and sham group.Mean±SD,*p<0.05,**p<0.01,***p<0.0001 indicate significant differences,and ns>0.05 means no significant difference.

## Discussion

The inflammatory response is a key mechanism by which the heart responds to injury and develops adaptive remodeling^[24]^. However, uncontrolled inflammation can weaken the adaptive response and promote heart damage. Both diabetes and hypertension-induced stress overload on the heart can lead to chronic inflammation and activation of macrophages^[25]^. Although macrophages play a crucial role in heart remodeling, the imbalance between pro-inflammatory and anti-inflammatory macrophage phenotypes can promote excessive inflammation to cause heart damage^[26]^. Myocardial fibrosis is an inevitable consequence of chronic myocardial injury^[27]^. On one hand, the activation and phenotype control of fibroblasts are heavily dependent on the inflammatory environment^[28]^. On the other hand, fibroblasts can also influence the macrophage phenotype. For example, in fibrosis and cancer, fibroblasts can support macrophages by providing CSF1, which can promote the survival of macrophages or facilitating signaling of macrophage-specific transcription programs. During inflammation, fibroblasts continue to provide CSF1 as a necessary factor for macrophages^29,30^.

The underlying mechanism of the interaction between fibroblasts and macrophages is highly complex, and they play important roles in various disease processes^[31]^. In our study, we found that there are considerable interactions between cardiac fibroblasts and monocyte/macrophages in TAC surgery-induced cardiac pressure overload, and PTN has a unique signal specificity. In addition, of all cell types, only cardiac fibroblasts express *Ptn*, and it was upregulated in TAC group compared with control group. Moreover, Under hypoxic condition *in vitro*, cardiac fibroblasts could secrete more PTN with the duration of hypoxia. In fact, *Ptn* is involved in a large number of biological processes. Its high expression in various cancers makes it an excellent biomarker and a target for anticancer drug development^[32]^ for gastric cancer^[33]^, lung cancer^34,35^, and glioblastoma^36,37^. PTN is associated with many important physiological events, including the maintenance of hematopoietic stem cells, adipocyte differentiation, chondrocyte development, and endothelial cell migration^[38]^. In recent years, some clues about PTN had also been found in the cardiovascular field. Studies had shown that PTN could enhance the survival of cardiomyocytes through inhibiting endogenous AKT/PKB activity^[39]^. PTN was also considered to be a potential biomarker for inflammatory ischemic cardiomyopathy with good ICM recognition^[40]^. In a proteomic analysis of pleural effusion from dead COVID-19 patients, 182 protein biomarkers including PTN were significantly elevate, indicating immune system overstimulation and cytokine storm^[41]^. These studies suggested that PTN was involved in the development of some cardiovascular diseases, but its underlying mechanisms are not well understood.

SDC4, one of the PTN ligands, has been found in recent years to be a biomarker for HF and myocardial injury^[42]^. SDC4 is significantly correlated with left ventricular geometric parameters in chronic heart failure, and serum SDC4 levels were associated with left ventricular size^[43]^. SDC4 is a proteoglycan with extracellular heparin sulfate chains that served as a central mediator for cell adhesion, migration, proliferation, endocytosis, and mechanical transduction. Its wide range of effects are reflected in its unique versatility in extracellular, membrane, and intracellular interactions. SDC4 was initially described as a ubiquitous low-affinity co-receptor for heparin-binding growth factors, but it is now thought to independently control numerous extracellular and intracellular signaling processes^[44]^. The broad function of SDC4 is partly due to its heterogeneity in ligand-binding capacity and partly due to its ability to interact with many intracellular signaling partners^[45]^. For example, SDC4 could promote the activation of PKCα, which in turn affect mTORC2 assembly^[46]^. SDC4 could also stabilize the interactions between growth factors and other cell membrane receptors.

FGF could bind to FGFR with high affinity, but this interaction and subsequent signaling events are amplified by the presence of heparin sulfate chains^47,48^. In this study, we found that *Sdc4* was widely expressed and abundant in various macrophage clusters as well as in fibroblasts (only one cluster) of TAC samples, and WGCNA analysis showed that the module containing the receptor SDC4 on monocytes/macrophages was significantly associated with the phenotype. This finding suggested that *Sdc4* may be one of the key genes involved in the cardiac stress overload process. The gene function of this module was mainly enriched in cytokines and inflammation-related genes, indicating that SDC4 had a similar effect on the biological functions of macrophages and indirectly suggesting the role of PTN on macrophages. To validate this hypothesis, we treated mouse macrophages Raw264.7 with recombinant mouse PTN, and found that the expressions of *TNF-α*, *Il6*, and *Cox2* were upregulated as well as the expressions of M2-type macrophage marker genes such as *Egr2*, *IL10*, and *Cd83* were downregulated^[49]^. Although we did not explore the entire expression profile of macrophages after PTN treatment, the current data suggested that PTN had an effect on the polarization of macrophages. Although the characteristics of macrophages *in vivo* are complex and variable across different phenotypes^50,51^, *in vitro* models provide a snapshot of the extreme phenotypes. In addition, TAC mouse model with myocardial hypertrophy was successfully established in our study, and we found that the the fluorescence intensity ratio of PTN/vimentin was significantly higher in TAC group compared with sham group, indicating an increased expression of PTN in fibroblasts in TAC group.

The various cells in the heart have a complex autocrine and paracrine network. The autocrine behavior of cardiac fibroblasts is more prominent under pressure overload^[14]^. The FGF2-FGFR2 ligand-receptor pair has been shown to regulate the release of pro-hypertrophic factors in fibroblasts^[52]^, and cardiac fibroblasts can release CCN2, which acts on multiple receptors including integrin receptors, heparin sulfate proteoglycans, and other receptors, and is an important autocrine-promoting fibrosis loop in cardiac fibrosis^[53]^. Fibroblasts can also secrete CGRP to induce anti-fibrotic effects^[54]^.

Certainly, there are some limitations to our research. Firstly, we lack experimental content regarding PTN gene knockout, therefore further investigation is required to understand the biological functions of PTN in macrophages and fibroblasts. Secondly, macrophage polarization requires a wide range of indicators, and we lack whole-transcriptome sequencing data, thus unable to provide a comprehensive description of the impact of PTN on macrophage expression profiles. Therefore, the biological functions and mechanisms of the PTN pathway still require further exploration.

## Authors contributions

Ke Sheng: Data curation, Investigation, Methodology, Writing-original draft

Yuqing Ran: Methodology

Yuting Guan: Methodology

Pingping Tan: Methodology

Rongrong Zhang: Methodology

Songwei Qian: Methodology

Hongzhou Lin: Methodology

Huilan Wu: Methodology

Yongmiao Peng: Methodology

Yuqing Huang: Methodology

Zhiguang Zhao: Methodology

Guanghui Zhu: Methodology, Investigation

Weiping Ji: Methodology, Investigation

Xiaoling Guo: Investigation, Conceptualization, Funding acquisition, Project administration, Writing-review & editing

## Acknowledgements

We thank all technicians of Basic Medical Research Center, the Second Affiliated Hospital and Yuying Children’s Hospital of Wenzhou Medical University.

## Data availability

Data will be made available on request.

---

**Fig. S1. The violin plots display the distribution of various characteristics in the control cells, such as feature counts, the proportion of mitochondrial RNA in each cell, and the proportion of hemoglobin RNA.**

**Fig. S2. The expression level distribution of mitochondrial RNA, nFeature_RNA, and hemoglobin RNA in individual cells of the control group as well as the proportion of outlier cells within the total population.**

**Fig. S3. Violin plots represent the distribution of different characteristics in the cells of TAC group, including feature counts, the percentage of mitochondrial RNA in each cell, and the percentage of hemoglobin RNA.**

**Fig. S4. The scatter plot reflects the expression level distribution of mitochondrial RNA, nFeature_RNA, and hemoglobin RNA in individual cells of the control group as well as the proportion of outlier cells within the total population.**

**Fig. S5. Moncle3 analysis of fibroblasts in the TAC group, depicted as a curve that transitions from the left starting point to the surrounding midpoint.**

**Fig. S6. Expression level variations of different ligand receptor members within the PTN pathway in fibroblasts of the TAC group as a function of simulated time.**

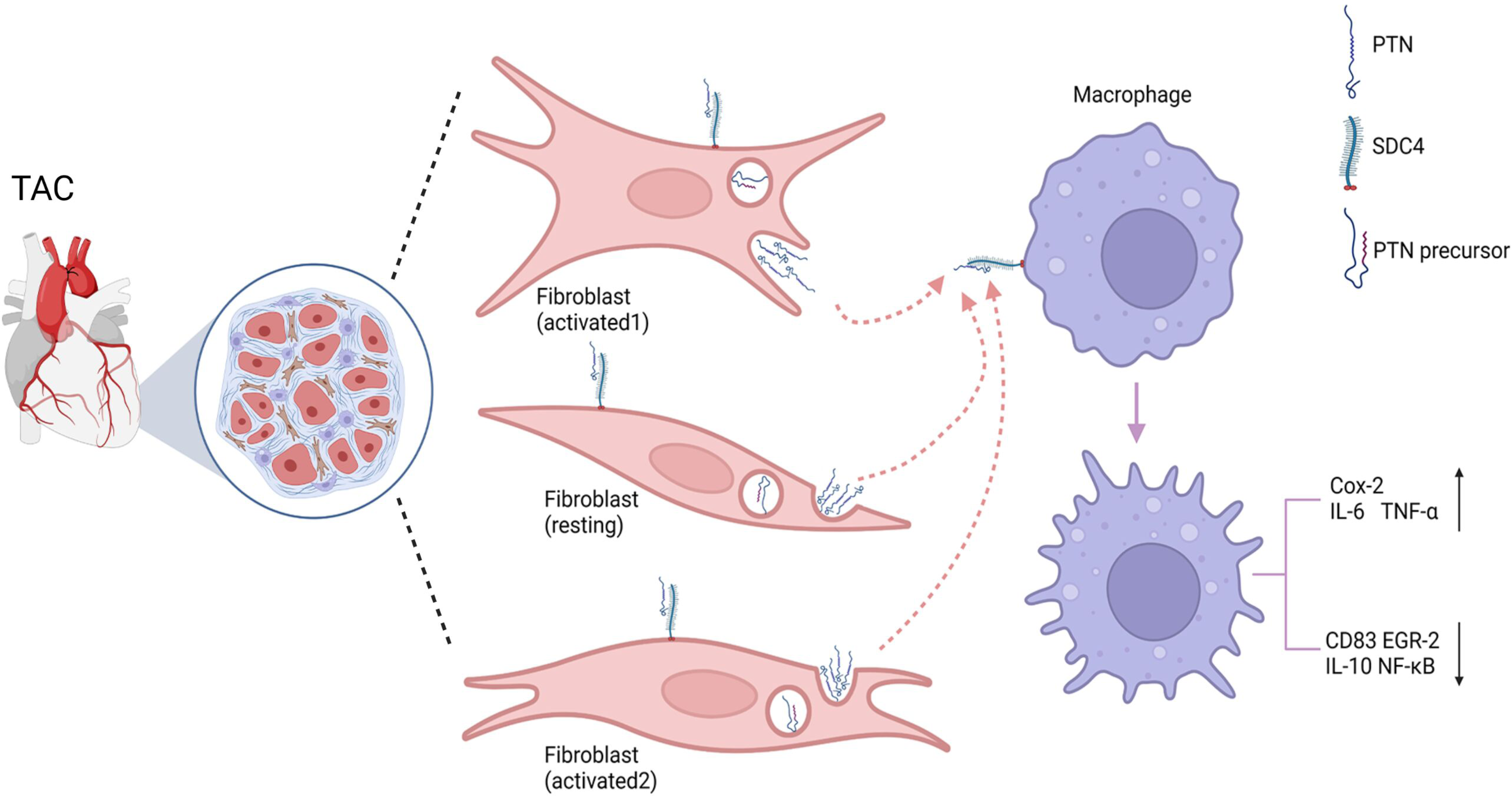

